# Male brain processing of the body odor of ovulating women compared to that of pregnant women

**DOI:** 10.1101/2020.10.16.340463

**Authors:** Ute Habel, Christina Regenbogen, Catharina Kammann, Susanne Stickel, Natalia Chechko

## Abstract

Female chemical signals underlie the advertising of sexual receptivity and fertility. Whether the body odor of a pregnant woman also has a signaling function with respect to male behavior is yet to be conclusively established. This study examines how the body odors of ovulating and pregnant women differentially affect the behavior of heterosexual men.

Body odor samples were collected from 5 pregnant women and 5 matched controls during ovulation. In a double-blind functional magnetic resonance imaging design, 18 heterosexual men were exposed to female body odors during ovulation (OV) and pregnancy (PRG) while being required to indicate the attractiveness of concurrently presented female portrait images. The participants were also required to indicate whether they assumed a depicted woman was pregnant.

While neither OV nor PRG altered the perceived attractiveness of a presented face, the men tended to identify the women as pregnant while exposed to a PRG body odor. On the neural level, OV activated a network of the frontotemporal and limbic regions, while PRG activated the superior medial frontal gyrus.

The results suggest that the detection of sexual availability activates the male brain regions associated with face processing and reward/motivation, whereas sensing pregnancy activates a region responsible for empathy and prosocial behavior. Thus, the female body odor during pregnancy likely helps foster circumstances conducive to the future care of offspring while the body odor advertising sexual availability promotes mating behavior. The brains of heterosexual men may be capable of unconsciously discriminating between these two types of olfactory stimuli.

## 1 Introduction

In nonhuman mammals, changes in the female scent are thought to mediate communication between the sexes, primarily facilitating mate choice (Mitchell et al., 2017) and optimizing and synchronizing reproductive activity (Coombes et al., 2018; Crawford and Drea, 2015). There is growing evidence that human body odor likely communicates similar signals (Lundström et al., 2008), mediated (analogous to studies in mammals (Takahashi, 1990)) by the hormonal changes across the female menstrual cycle. Several studies have demonstrated that the body odor of (near-)ovulating women is rated as more pleasant and sexually arousing by heterosexual men compared to that of the same women in their luteal phase of the menstrual cycle (Gildersleeve et al., 2012; Havlicek et al., 2006; Singh and Bronstad, 2001). Lobmaier et al. (2018) have observed a male preference for the body odor of women with higher estradiol and lower progesterone levels, which represent the late follicular phase, leading to ovulation. The use of oral contraceptives, on the other hand, has been found to result in an absence of body odor preference in men as the contraception prevents ovulation (Kuukasjarvi et al., 2004; Miller et al., 2007). Similar to the changes in female body odor during ovulation, facial characteristics are also modulated by cyclic hormonal changes. In fact, photographs of women taken during their most fertile days have been rated more attractive than those taken during the luteal phase (Puts et al., 2013; Roberts et al., 2004). In heterosexual men, increased testosterone levels in response to female scents (Cerda-Molina et al., 2013; Miller and Maner, 2010) may help, as indicated by studies in nonhuman mammals, increase male mating rates though enhanced spermatogenesis or production of attractive male chemosignals (for a review see Coombes et al. Coombes et al., 2018). As ovulation indicates the best time for reproduction, it appears that men are capable of unconsciously distinguishing between ovulating and non-ovulating women, with a preference for the former. With most research attention having been directed, even in human studies, to the function of female chemical signaling in the context of sexual receptivity and fertility, signals outside of fertility (e.g., during pregnancy) have not been adequately explored. Studies exploring nonhuman mammals have offered tentative evidence of reduced sexual advertisement during pregnancy (for a review see Coombes et al. Coombes et al., 2018). Male golden hamsters, for instance, have been found to be less lured by the scents of pregnant females (Johnston, 1980). Pregnant or lactating female hamsters seem to have the lowest scent marking rate in these phases (Johnston, 1979), indicating reduced sexual advertising to avoid male advances. Reduced male sexual desire in response to pregnant or lactating females also likely facilitates social behavior that ensures the protection of pregnant females and their offspring. The olfactory changes in pregnant and lactating rodents are linked to their sexual non-availability and, thus, the protection of their offspring (Coombes et al., 2018). The male banded mongoose has also been found to spend significantly more time exploring non-pregnant females compared to pregnant females (Mitchell et al., 2017).

Given that several hormones, including estradiol, progesterone and human chorionic gonadotropin (HCG), increase dramatically during pregnancy (Tulchinsky et al., 1972), the hormonal composition deviating from the normal menstrual cycle should have an influence on body odor. Indeed, olfactory samples of the para-axillary lines and the nipple regions during pregnancy have been found to consist of five different components, which dissipate following pregnancy (Vaglio et al., 2009).

To date, there has been no investigation of the neural network underlying the processing of female body odor during pregnancy in comparison to ovulation. This study seeks to explore the extent to which ovulation and pregnancy in humans may be recognized through olfactory signals and how they influence behavior and the underlying brain activation in heterosexual men. In previous brain imaging studies, the neural signature of face processing in general has been found to comprise a widespread network of the occipitotemporal cortex, with the judgment of facial attractiveness additionally recruiting an extended system of the limbic (the amygdala, the hippocampus, the cingulum, the anterior insula) and reward-related structures (the orbitofrontal regions, the amygdala, the basal ganglia) (for a review, see Hahn et al. Hahn and Perrett, 2014). Thus, we expected that the odor of ovulating women (OV), compared to that of pregnant women (PRG), would result in higher facial attractiveness ratings (Hypothesis I) and trigger neural responsivity in areas related to the perception of female attractiveness in heterosexual men (Hypothesis II). Additionally, we explored whether there may be an underlying neural responsivity to the unconscious discrimination between OV and PRG in men as those with the ability to make this distinction likely have an advantage when it comes to sexual selection. We expected that the simultaneous presentation of PRG and the photographs would bias the male participants toward identifying more women as pregnant compared to the presentation of OV (Hypothesis III) and that, during pregnancy categorization, PRG would prompt a stronger recruitment of the brain network responsible for facial identity (Hypothesis IV).

## 2 Materials and Methods

### 2.1 Participants

The study involved two separate groups of participants: odor donors and odor recipients. Please see the SI Material and Methods for detailed information on the recruitment, dietary restrictions and odor sampling procedure.

Odor donors: 5 healthy, non-pregnant, ovulating Caucasian women with a steady natural menstrual cycle and not using any form of contraception (age: range = 21 – 30 years; M = 24 years, SD = 3.96) and 5 healthy pregnant women in their first trimester of pregnancy (age: range = 24 – 36; M = 30.2, SD = 4.47) were recruited. The exclusion criteria for both groups included smoking, taking any type of medication, and legal or illegal drug abuse. Two body odor pads (left and right armpit) were obtained from each donor on each of the two nights of participation. In total, there were 20 pads from pregnant and 20 pads from ovulating women.

Odor recipients: 20 healthy, Caucasian, single, right-handed, heterosexual male (age: range = 20 – 29 years, M = 24.74, SD = 2.92) participants were recruited at the RTWH University Aachen. The exclusion criteria included deviant sexual orientation, being in a relationship, smoking or consuming any type of legal or illegal drugs, any type of medication, frequent nosebleed or nosebleed three days prior to recruitment, current infections or diseases of the nose or the respiratory system, and body mass index below or above the normal range (BMI normal range for men aged 20-30 years: 20-26). To control for depression or any type of psychiatric disorders, BDIII (Hautzinger et al., 2006) and SCID light (Structured Clinical Interview for DSM-IV; Wittchen et al., 1997) were performed by an experienced psychologist (SS). Olfactory performance was assessed using the Sniffin’ Sticks identification test (Hummel et al., 1997).

One participant did not follow the dietary restrictions, and one participant did not appear. Since thawing of the body odor pads took place 30 minutes before the functional magnetic resonance imaging (fMRI) experiment, one OV pad and one PRG pad were rendered unusable. In total, the data of 18 male participants were included in the analyses.

Written informed consent was obtained from all participants. The study protocol was in accordance with the Declaration of Helsinki and approved by the Institutional Review Board of the Medical Faculty, RWTH Aachen University.

### 2.2 Procedure

This study was conducted in two different phases (Figure 1). First, the female odor donors were recruited and screened, and the odor and blood samples were collected and frozen. Second, the male participants were recruited and exposed to the odor samples while being examined in the fMRI. For further details, please see SI Material and Methods.

**Figure 1.**
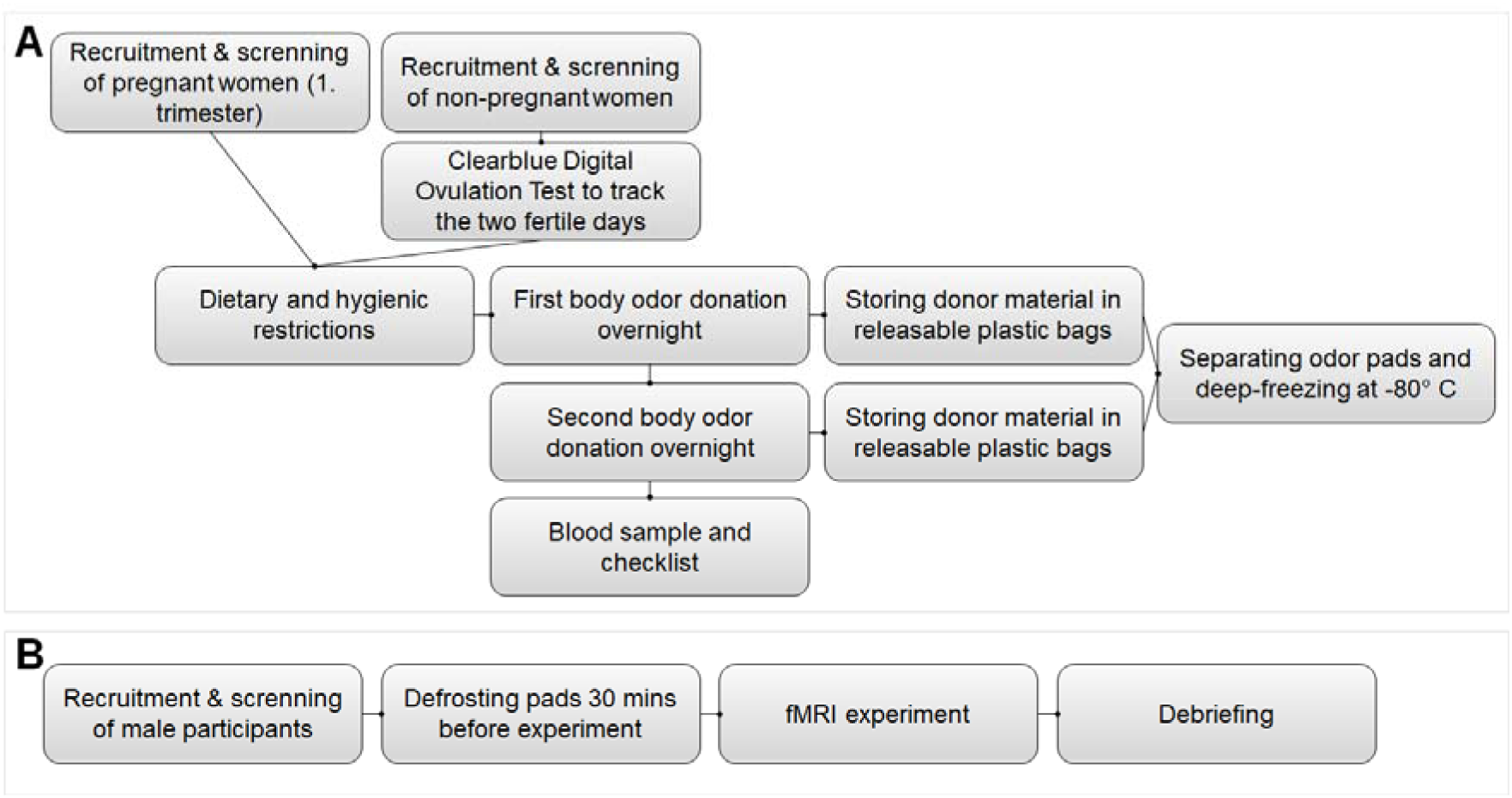
A) Odor donors: Collection of women’s body odors. B) Oder recipients: Male testing procedure.

### 2.3 Experimental task

#### 2.3.1 Visual and olfactory stimulation

The stimuli comprised combinations of olfactory and visual stimuli. Standardized portraits of emotionally-neutral female faces (Oslo Face Database, Chelnokova et al., 2014) were used for visual stimulation (see also SI Material and Methods).

The olfactory stimuli were body odor samples of the female odor donors (ovulation and pregnancy). An unused pad containing no odor was used as a no-odor (NO) condition, which had also been deep-frozen at −80° Celsius.

For each run in the fMRI experiments (each experiment consisted of three runs), body odor pads of one odor condition (either PRG, OV, or no NO) were fixated under the participants’ nose using an odorless Leukotape. Having been informed that the stimulus pads might or might not smell of anything, and that they should not pay attention to the smell but concentrate on the faces, the participants were not aware that they would be exposed to body odor. After each odor condition, the participants were taken out of the MRI unit for the removal of the used pad and the attachment of another for the next odor condition.

The presentation of the visual stimuli during the fMRI tasks and the recording of the subjects’ feedback and scanner triggers were achieved using the software E-Prime 2.0 (Schneider et al., 2008). The presentation was projected onto an MRI-compatible screen, visible via a mirror mounted to the head coil. In addition to oral instructions prior to the measurement, all participants read the instructions on the screen in the MRI scanner (see also SI Material and Methods).

#### 2.3.2 Tasks

Two experiments (attractiveness rating and pregnancy categorization) with three runs each were conducted representing three different odor conditions (PRG, OV, NO). During each odor condition, 10 different photographs were drawn from a chosen pool of 30 images (Chelnokova et al., 2014) which were presented twice in a randomized order for each olfactory condition and participant to ensure that neither the individual photograph nor the order of photographs could have a confounding effect. Both fMRI experiments were presented in a mixed block and random order.

##### Attractiveness rating: Effect of chemosensory stimulation on subjective perception of attractiveness (Figure 2 A)

**Figure 2.**
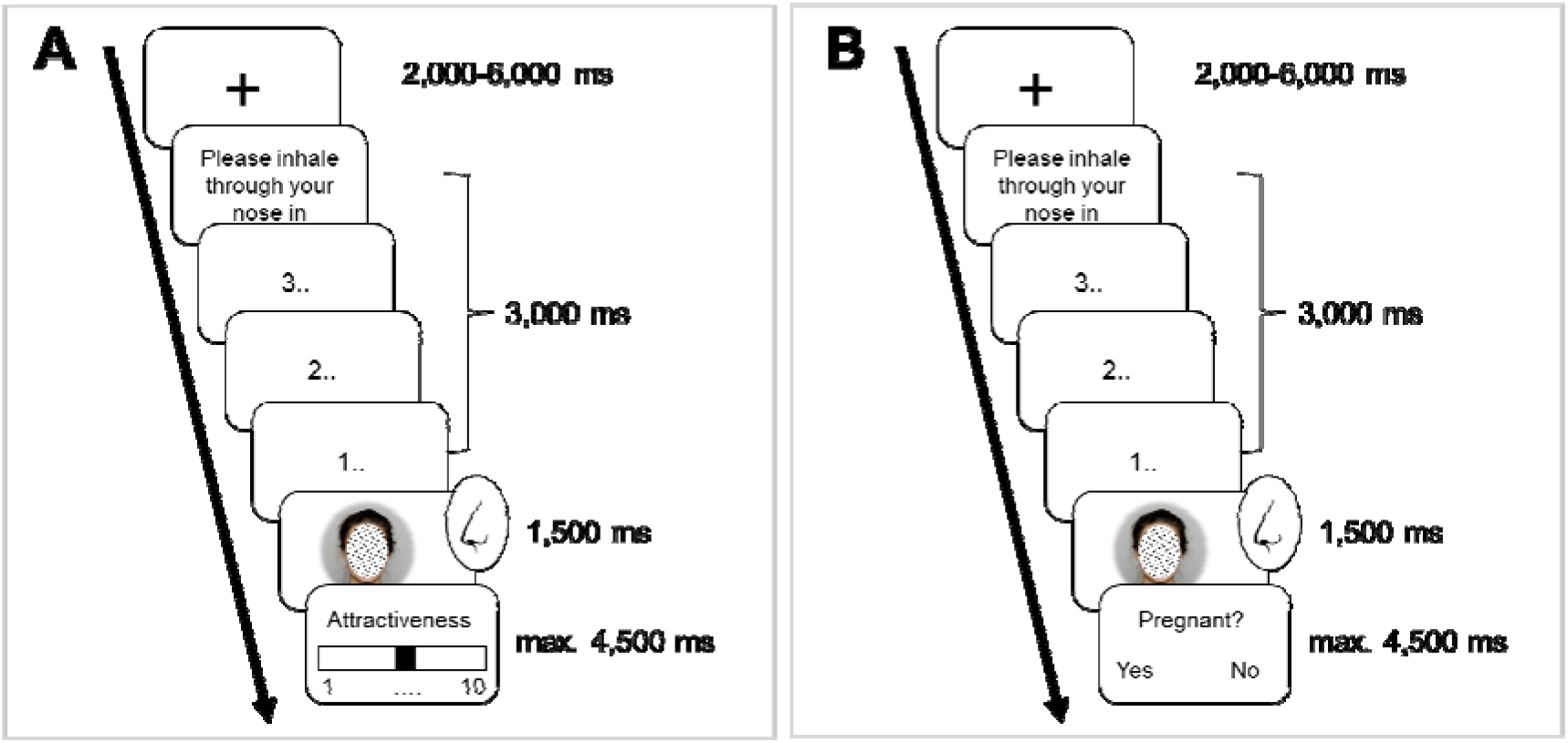
Experimental procedure of the functional tasks. A) Attractiveness rating: Unconscious perception of ovulation and pregnancy. B) Pregnancy categorization: Conscious perception of ovulation and pregnancy.

Each trial started with a black fixation cross (jittered 2,000 – 6,000 ms) in the center of a white screen. To ensure that participants would inhale through their nose while being exposed to each photograph, a countdown before each image presentation was used as follows: “Please inhale through your nose in 3..2..1” (total 3,500 ms). As the participants inhaled, a face was presented for 1,500 ms. After each image, participants rated the attractiveness of the woman portrayed using the LUMItouch response system (LUMItouch, Photon Control, Burnaby, Canada), a button press device held in the right hand. The index and ring fingers were used to scroll from 1 = not attractive at all to 10 = very attractive and the middle finger to confirm the entry.

##### Pregnancy categorization: Effect of chemosensory stimulation on subjective perception of pregnancy (Figure 2B)

The images were presented in the same manner as described above. This time, participants were asked directly to indicate whether or not they thought the depicted person was pregnant. They were asked to respond using the index finger for “yes” and the middle finger for “no”.

#### 2.4 fMRI data acquisition and analysis

The neuroimaging data were acquired using a 3T Trio Prisma MR scanner (Siemens Medical Systems, Erlangen, Germany) located in the Medical Faculty of the RWTH University Hospital in Aachen. Functional images were collected with an echo-planar imaging (EPI) T2* weighted contrast sequence which is sensitive to blood oxygen-level-dependent (BOLD) contrast (voxel size: 3 × 3 × 3 mm^3^, 64 × 64 matrix, FoV: 192 × 192 mm^2^, 34 slices, whole-brain acquisition, descending, no spacing between slices, TR = 2 s, TE = 28 ms, alpha = 77°). On average, each of the six EPI sequences took 5 minutes with approximately 120 scans. High resolution, T1-weighted structural images were acquired by means of a three-dimensional MPRAGE sequence (voxel size: 1 × 1 × 1 mm^3^, sagittal FoV: 256 × 256 mm^2^, 176 slices, TR = 5.0 s, TE = 2.98 ms, alpha = 4°). The duration of the MPRAGE sequence was 8:22 min.

The imaging data were pre-processed and analyzed using Statistical Parametric Mapping 12 (SPM12; https://www.fil.ion.ucl.ac.uk/spm/) implemented in Matlab 2015b (Mathworks Inc., Natrick, Massachusetts, United States). The first three images of each time series were discarded due to T1 stabilization effects. Preprocessing involved the adjustment of the origin of all images to the anterior commissure before realignment. Functional scans were spatially corrected for individual head movements during the realignment step and co-registered to the anatomical scan. The anatomical image was segmented in white and gray matter and the cerebrospinal fluid with tissue probability maps. Approximations of spatial normalization parameters in the MNI standard space were calculated (voxel size = 3 × 3 × 3 mm^3^). These parameters were adapted to the functional images and used in the normalization step. The normalized EPI data were spatially smoothed with an isotropic Gaussian kernel (full-width-at-half-maximum = 8 mm). All coordinates are in reference to the Montreal Neurological Institute (MNI) convention (http://www.mni.mcgill.ca).

For each subject, delta functions with the time points of each type of trial presentation were convolved with the canonical hemodynamic response function (HRF) to build a regression model of the time series. Realignment parameters including percent signal changes of each participant were included as nuisance variables. Supplementary HRF-convolved regressors of no interest were the onsets of the rating scales, instructions and countdowns. A high-pass filter with a cut-off period of 128 s was applied and serial auto-correlations were accounted for by including a first-order autoregressive covariance structure (AR(1)). At individual levels, for each task (attractiveness rating and pregnancy categorization) a simple main contrast was computed for OV, PRG, and NO.

An SPM12 random-effects analysis was performed by entering all six contrasts into a mixed-effects design for second level group analyses. To correct for multiple comparisons, when not otherwise mentioned, we applied extent threshold correction as defined by Monte Carlo simulations (3DClustSim; implemented in AFNI; Cox, 1996) to prevent false-discovery rates. For a threshold at the voxel level of p = .001 uncorrected, and spatial properties of the current study, 10,000 iteration resulted in an extent threshold of k = 67 resampled voxels, corresponding to a cluster-level familywise error (FWE) of p < .05.

### 2.5 Analysis of Behavioral Data

The attractiveness ratings and pregnancy categorization collected during the functional imaging tasks were analyzed via IBM SPSS statistics 21. According to Hypothesis I, a repeated measures ANOVA should reveal a main effect of odor (OV, PRG, NO; within-subject factor) on the attractiveness rating with OV expected to yield the highest attractiveness ratings of the presented faces.

Using the generalized estimating equation (GEE) method with a Poisson loglinear model and an autoregressive covariance structure, we stated in Hypothesis II that the frequency of categorizing a female face as pregnant (dependent variable) would differ between the conditions (within-subject variable) across the participants (subject variable) with PRG expected to yield the highest frequency of categorizing faces as pregnant.

To check for order effects in attractiveness ratings and pregnancy categorization, post-hoc tests were employed with different combinations of the application order.

### 2.6 Blood Analysis

Blood samples from both female odor donor groups were taken on the days before the first and the second odor donations and examined at the laboratory diagnostic center of the RWTH Aachen University Hospital using the ECLIA (electrochemiluminescence) method to ensure that the hormonal concentration (estradiol, progesterone, HCG) pattern would be representative of the corresponding reproductive phase.

## 3 Results

### 3.1 Behavioral results

A repeated measures ANOVA revealed no main effect of odor (F(2, 34) = 1.456, p = .247) on attractiveness ratings, contrary to Hypothesis I (Figure 3A). Contrary to Hypothesis II, the GEE analysis revealed no significant main effect of condition (Wald-*χ*^2^(2) = 5.056, p = .08) on pregnancy categorization. On the descriptive level, the participants were found to rate faces during OV odor stimulation less often as pregnant compared to both PRG stimulation and a no-odor (NO) condition (see Figure 3B).

**Figure 3.**
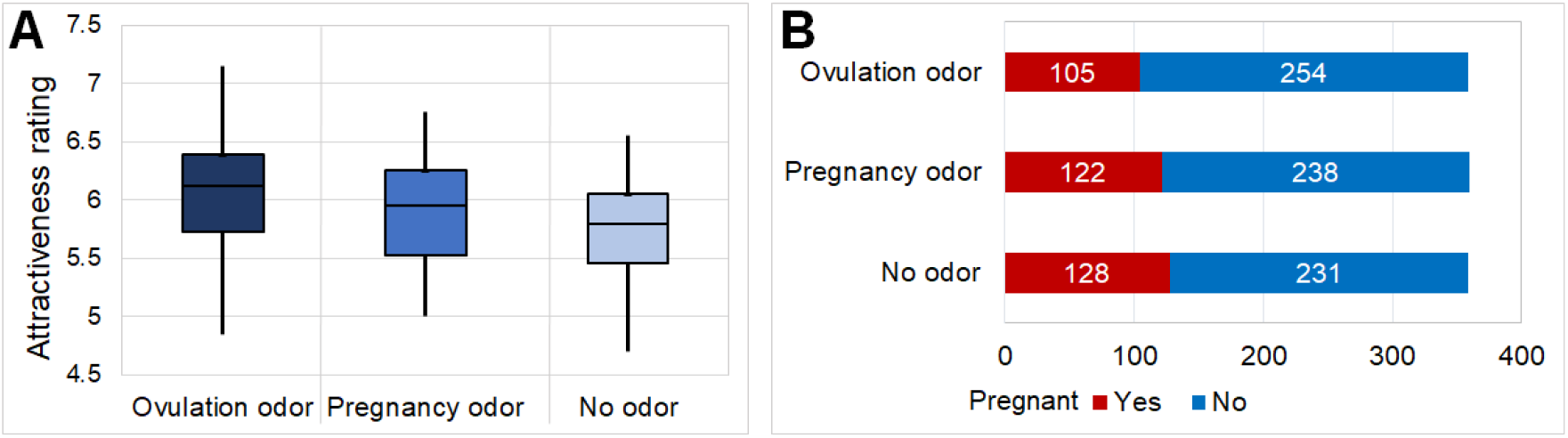
A) Box plots of mean attractiveness ratings OV: M = 6.04, SD = 0.65; PRG: M = 5.89, SD = 0.51; NO = 5.78, SD = 0.54), with 25^th^ and 75^th^ percentile as well as minimum and maximum values. B) Overall frequency of pregnancy categorization.

Post-hoc tests evaluating order effects revealed no significant differences between OV presented first vs. last (n = 11) either in attractiveness ratings (p = .191) or in pregnancy categorization (p > .188), and no significant differences between PRG presented first vs. last (n = 14; p = .707; p > .49) or NO presented first vs. last (n = 13; p = .721, p > .786).

The means, the standard deviations and ranges of estradiol and progesterone of the ovulating and pregnant odor donors as well as the HCG levels of the pregnant odor donors are presented in Table 1. Both estradiol and progesterone differed significantly between OV and PRG.

**Table 1.** Mean (M), standard deviation (SD) and range of ovarian hormone concentration in ovulating and pregnant odor donors.

|  | OV |  | PRG |  | t(df = 31) | P |
| --- | --- | --- | --- | --- | --- | --- |
|  | M (SD) | Range | M (SD) | Range |  |  |
| Estradiol (pg/ml) | 71.04 (43.07) | 24.30 – 155.20 | 2169.27 (1368.65) | 266.10 – 3996.00 | -6.32 | < .001 |
| Progesterone (ng/ml) | 2.57 (1.72) | 1.06 – 6.74 | 33.80 (14.29) | 18.03 – 56.92 | -8.94 | < .001 |
| HCG (mU/ml) | - | - | 63,073.75 (32,515.48) | 35,011.0 – 115,967.00 | - | - |

Note. OV: body odor collected during ovulation; PRG: body odor collected during first trimester of pregnancy.

### 3.2 fMRI results

#### Effects of body odor on whole-brain activation

In a whole-brain analysis, we explored whether neural processing was affected by the type of body odor. First, we sought to determine the activation pattern by contrasting OV against PRG and vice versa, independent of the task (attractiveness rating or pregnancy categorization) and for each task individually. The ovulation odor (OV > PRG) led to a significant activation of the right middle orbital gyrus (144 voxel, T = 4.65, MNI: 12/64/-8) and the left fusiform gyrus (67 voxel, T = 4.25, MNI: -28/-48/-12). No significant suprathreshold activation was observed in the pregnancy odor (PRG > OV).

#### Interaction effects of body odor and task

In the attractiveness rating task, ovulation odor (OV > PRG) revealed significant clusters in the left inferior frontal gyrus, the middle orbital gyrus, the left cerebellum, the right medial orbital and superior orbital gyrus, the left precuneus and calcarine gyrus, the right insula, the temporal pole and the amygdala, and the left supramarginal gyrus (Figure 4, Table 2). The pregnancy odor (PRG > OV) on the other hand did not yield significant activation clusters.

**Table 2.**
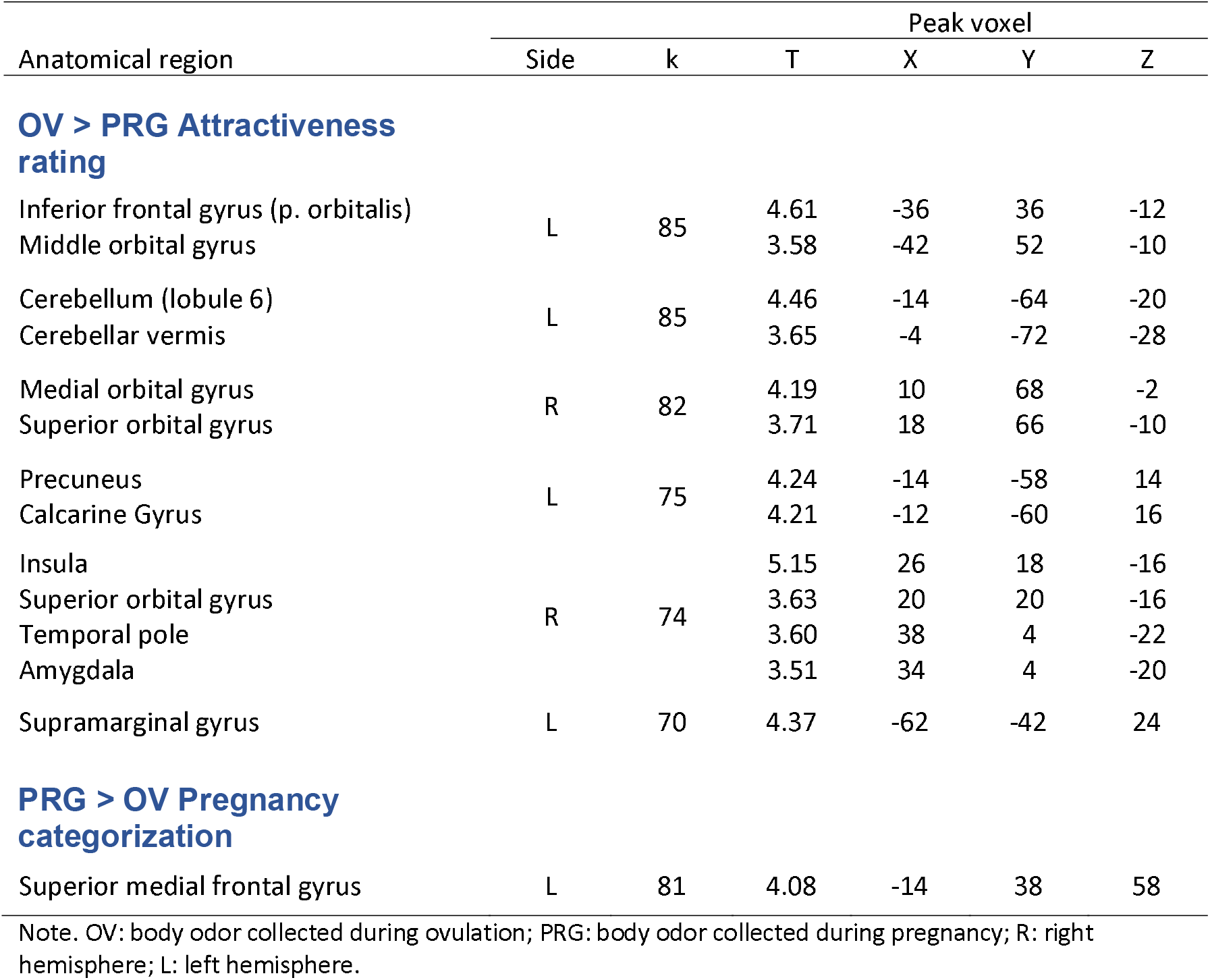
Effects of condition and task on whole-brain activation, with p < .001, with extend threshold k = 67 (corresponding to Monte Carlo correction)

| Anatomical region | Side | k | T | Peak voxel |  |  |
| --- | --- | --- | --- | --- | --- | --- |
|  |  |  |  | X | Y | Z |
| OV > PRG Attractiveness rating |  |  |  |  |  |  |
| Inferior frontal gyrus (p. orbitalis) | L | 85 | 4.61 | -36 | 36 | -12 |
| Middle orbital gyrus |  |  | 3.58 | -42 | 52 | -10 |
| Cerebellum (lobule 6) | L | 85 | 4.46 | -14 | -64 | -20 |
| Cerebellar vermis |  |  | 3.65 | -4 | -72 | -28 |
| Medial orbital gyrus | R | 82 | 4.19 | 10 | 68 | -2 |
| Superior orbital gyrus |  |  | 3.71 | 18 | 66 | -10 |
| Precuneus | L | 75 | 4.24 | -14 | -58 | 14 |
| Calcarine Gyrus |  |  | 4.21 | -12 | -60 | 16 |
| Insula | R | 74 | 5.15 | 26 | 18 | -16 |
| Superior orbital gyrus |  |  | 3.63 | 20 | 20 | -16 |
| Temporal pole |  |  | 3.60 | 38 | 4 | -22 |
| Amygdala |  |  | 3.51 | 34 | 4 | -20 |
| Supramarginal gyrus | L | 70 | 4.37 | -62 | -42 | 24 |
| PRG > OV Pregnancy categorization |  |  |  |  |  |  |
| Superior medial frontal gyrus | L | 81 | 4.08 | -14 | 38 | 58 |

Note. OV: body odor collected during ovulation; PRG: body odor collected during pregnancy; R: right hemisphere; L: left hemisphere.

**Figure 4.**
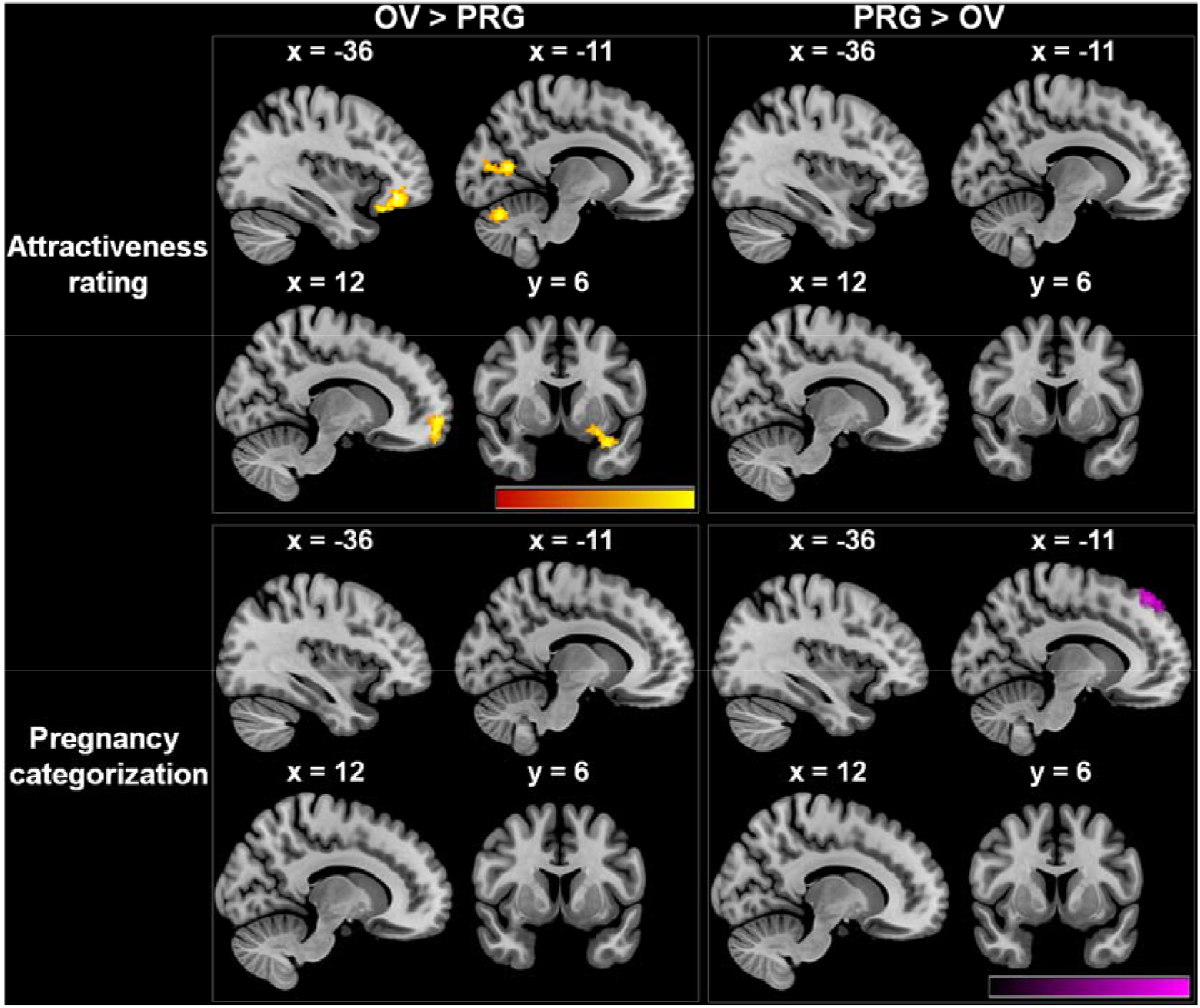
Four field matrix of brain regions activated during ovulation odor (OV) > pregnancy odor (PRG) and PRG > OV across both tasks. For visualization purposes, significant clusters are presented at an uncorrected threshold of p < .01.

In the pregnancy categorization task, no significant suprathreshold activation was observed for the ovulation odor (OV > PRG), whereas the pregnancy odor (PRG > OV) led to an increased activation of the left superior medial frontal gyrus (Figure 4, Table 2).

Please see the supplementary information (SI fMRI Results) for 1) analyses of the task effects (attractiveness rating versus pregnancy categorization and vice versa) independent of odor condition on whole-brain activation, and 2) for analyses concerning the effects of body odor compared to NO and vice versa on whole-brain activation.

## 4 Discussion

The current study investigated the behavioral and neural effects of female body odor exuded during two distinct reproductive stages, ovulation and pregnancy, on healthy, single, heterosexual men. While no clear effect of body odor was detected on the behavioral level, we observed distinct interaction effects, revealing differential neural activation patterns in response to the different odor conditions dependent of the task. Ovulation odor (OV > PRG) was found to lead to a crossmodal effect of increased neural activation in the orbitofrontal cortex and the left fusiform gyrus in response to visual stimuli. This was present in both tasks, but, specifically in attractiveness rating, ovulation odor was found to lead to a stronger involvement of the orbitofrontal cortex, the precuneus, the insula and also the temporal-limbic structures including the temporal pole and the amygdala. Thus, the body odor signaling sexual availability was found to lead to greater responsivity in the brain networks linked to the core system and the extended system involving face processing (the fusiform gyrus, the temporal pole, the insula and the amygdala) (Hahn and Perrett, 2014; Haxby et al., 2000) and the emotion- and reward-related regions responsible for esthetic judgments (the orbitofrontal regions, the amygdala) (Aharon et al., 2001; O’Doherty et al., 2003; Shen et al., 2016; Winston et al., 2007). Evidence suggests that attractive faces receive longer looks compared to unattractive ones (Leder et al., 2010) and therefore receive more attention as they lead to rewarding experiences (Hayden et al., 2007). Given that the chosen photographs were of average attractiveness, the mere odor of an ovulating woman appeared to crossmodally reinforce greater activity in a distributed neural network including the associative cortices, and the limbic and paralimbic areas. Apart from representing the processing of motivational and emotional information (Lang and Davis, 2006) (pleasant or appetitive stimuli tend to evoke greater attention and affective and physiological arousal), this brain network is also linked to sexual arousal evoked by visually presented erotic or pornographic stimuli (e.g. Arnow et al., 2002; Beauregard et al., 2001; Karama et al., 2002; Seok et al., 2016). Coupled with the fact that exposure to the smell of an ovulating woman can result in sexual desire (Cerda-Molina et al., 2013), the observed neural pattern in response to ovulation body odor may indicate that our study participants experienced greater attraction toward the depicted women and showed neural signs of (sexual) arousal.

Although human sexual behavior, unlike that of other species, is no longer solely confined to the purposes of reproduction, male sexual arousal in response to fertility cues may be an evolutionary remnant of a behavior that was necessary for the survival of the species. Given the analogous evidence of cerebral networks underlying attention, attraction and arousal in men in response to ovulation odor (OV > PRG), it can be assumed that the advertisement of receptivity with the aim of optimizing reproductive success, as seen in various nonhuman mammals (Coombes et al., 2018; Crawford and Drea, 2015), may also have been retained in humans. However, the absence of a corresponding behavioral effect suggests that the conscious perception of attractiveness is distinct from the chemosensory influence on the underlying neural correlates. An early study by Istvan et al. (1983) found sexually aroused participants (both male and female) not differing from their unaroused counterparts in terms of attractiveness ratings of (below average, average and highly attractive) female faces. Thus, while neither sexual arousal nor the chemosensory perception modifies perceived attractiveness, the neuronal processing of human odor, like sexual arousal, follows direct evolutionary mechanisms of reproduction.

Pregnancy odor (PRG > OV) was found to lead to a greater activation of the superior medial frontal gyrus in response to visual stimuli during the attractiveness rating task. In sexually motivated male rats, the medial prefrontal cortex is responsible for motivational processes during sexual behavior (Hernández-González et al., 2007) and lesions cause reduced initiation of sexual behavior (Ågmo et al., 1995).

Furthermore, studies in rodents suggest that the smell of a pregnant female can reduce sexual desire in males, resulting in them not approaching the female and her offspring (for a review, see Coombes et al. Coombes et al., 2018). In comparison to the neural pattern observed in response to the ovulation odor, the smell of pregnant women appeared to result in decreased facial processing and attention in our heterosexual male subjects. In humans, the dorsomedial prefrontal cortex, to which the medial frontal gyrus belongs, is a central part of the social brain network that enables people to understand and interact with one another (Frith and Frith, 2007; Seitz et al., 2006). Activity in the medial frontal cortex has been linked to empathy-related prosocial behavior, such as helping others (Rameson et al., 2012), and altruistic motivations (Basile et al., 2011; Moll et al., 2007). This region has been repeatedly identified as an appraisal dimension that facilitates social knowledge involving inferences of other’s internal thoughts and their evaluation (for a review see Wagner et al., 2012). The dorsomedial prefrontal cortex is functionally involved when an individual reflects on unobservable mental states of others, known as social person knowledge (Dixon et al., 2017; Wagner et al., 2012), or predicts them under uncertainty (for a review see Isoda and Noritake, 2013). This evaluation is made particularly with regard to outcomes that can potentially affect one’s well-being and either facilitate or hinder one’s goals (Isoda and Noritake, 2013).

Based on the results of our study, we postulate that the mere smell of a pregnant woman has an impact on a man’s evaluation of the opposite gender, which is also modulated by his own goals and intentions. This can either imply that empathic and prosocial brain networks in men are activated by the smell of pregnancy odor, or that this odor interferes with their personal goals and thus triggers stronger evaluation processes, reflecting the relevance of the particular situation compared to other conditions. To support these theories, we would of course need more data/results focusing on empathy-related factors, trustworthiness and the perception of vulnerability of pregnant women rather than their attractiveness.

In summary, we found distinct neural activation patterns in response to different body odors as well as an effect of the specific task. The smell of an ovulating woman had no significant effect on brain activation when the participants were required to determine whether or not someone was pregnant, nor did the smell of a pregnant woman when they rated the women’s attractiveness. Thus, correctly identifying a female as bearing offspring or being attracted to one likely serves greater evolutionarily-relevant purposes. In other words, it seems that body odors not only chemically signal someone’s reproductive status, but also prepare the receiver to instinctively initiate appropriate cognitive processes. Consequently, the widespread activations seen during the NO condition (compared to OV as well as PRG; see SI fMRI Results) may simply reflect the absence of crossmodal stimulation. The absence of reproductive chemosensory signals prompted the participants to stringently perform only two unimodal tasks: to assess the attractiveness of a female face and to categorize her as pregnant or not pregnant.

### Strengths, limitations and conclusions

This, to our knowledge, is the first study to examine the effects of body odor of ovulating and pregnant women on male heterosexuals both in terms of behavior and brain activity. Previous studies have been mutually exclusive by focusing on either visual attractiveness during the menstrual cycle using photographs or body movements (Miller et al., 2007; Puts et al., 2013; Roberts et al., 2004), or on the attractiveness of body odor over the entire cycle (Cerda-Molina et al., 2013; Gildersleeve et al., 2012; Havlicek et al., 2006; Kuukasjarvi et al., 2004; Lobmaier et al., 2018; Probst et al., 2017). Our attempt is the first to combine the investigation of both the neural and behavioral effects of reproductive body odor on facial attractiveness and pregnancy ratings. However, we did not present the body odor of a woman along with her own photograph, but used random female faces with average attractiveness. While other studies have used a wider variety of faces (ranging from attractive to unattractive), thus showing differences in the respective task requirements (e.g. Foo et al., 2017; Lucas and Koff, 2013), we chose not to focus on the basal differences in attractiveness when selecting averagely attractive women. We conclude, therefore, that the body odors of OV and PRG had no influence on the attractiveness ratings of averagely attractive women’s faces. However, the two types of body odor triggered different patterns of activation: while the body odor of ovulating women activated the areas associated with face processing as well as reward- and motivation-related processing, the pregnancy scent did not.

The following limitations should also be considered in the context of our findings. The group sizes of the female odor donors and the male participants were relatively small. Since on average 5 male participants smelled the very same body odor, the individual measurements were not independent of one another. To allow for a greater variance, odor samples should include mixed odors of several donors. Also, the male participants were not asked how pleasant or attractive they found the individual smells or whether they were sexually aroused. These and other details (such as other perceived aspects of the face, apart from attractiveness) could provide further insight into the basal effects of female body odor.

In sum, similar to what has already been observed in animal studies (Coombes et al., 2018; Crawford and Drea, 2015), we can assume that women’s body odor not only changes within the menstrual cycle, but it also takes on other characteristics during pregnancy. In view of this, signaling sexual unavailability (as described in primates and non-primates (Crawford and Drea, 2015; Johnston, 1980; Mitchell et al., 2017)) may also be important in humans. Based on our observation and the findings of animal studies, we assume that the different activity patterns during OV (facial processing, reward-/motivation-related processing) and PRG (social brain) elicit behavioral results that are beneficial, in evolutionary terms, not only to the man, in terms of mate choice, but also to the woman, as her pregnancy likely prevents sexual advertising and triggers prosocial behavior in men. Given the significantly disparate neural activations in OV compared to PRG, the female body odor in pregnancy may indeed change communications between genders.

## Acknowledgments

This study was supported by the International Research Training Group (IRTG 2150) of the German Research Foundation (DFG). The authors are grateful to the odor donors and fMRI participants for their time and effort.

## Author Contributions

UH, CR, SS, and NC: Conceptualization; SS: Data curation; SS: Formal analysis; CK and SS: Investigation; CR and SS: Methodology; UH and NC: Project administration; CR and SS: Software; UH and NC: Supervision; Writing - original draft: UH, SS, and NC; Writing - review & editing: UH, CR, SS and NC.

