## Supplementary Information for "Male brain processing of the body odor of ovulating women compared to that of pregnant women"

### **SI Materials and Methods**

### **Procedure**

Phase 1: Collection of Women’s Body Odors

Five non-pregnant women not using any hormonal contraception were recruited at the University Hospital RWTH Aachen. They were asked to collect body odor using armpit pads during their ovulation phase. They were instructed to use the Clearblue Digital Ovulation Test every morning of their menstrual cycle until the ovulation test indicated an increase in the concentration of the luteinizing hormone (LH). They could use the cycle calculator provided by Clearblue for orientation. The advantage of this method is that even those with a very short cycle can correctly determine their time of ovulation. A rise in LH concentration indicates the beginning of the two most fertile days of the menstrual cycle (Ecochard et al., 2001). At this point, the non-pregnant participants were asked to make an olfactory donation on both days.

Five women during their first trimester of pregnancy (8 to 14 weeks) were recruited from the prenatal medicine department of the University Hospital RWTH Aachen as well as from registered gynecologists. Eligible women were asked to show their mother-child passport so that the week of pregnancy could be checked and noted.

On the day before the first and second odor donations, a blood sample was taken from all donors to carry out a hormone validation (determination of progesterone and estradiol). Additionally, HCG (Human chorionic gonadotropin) blood values of the pregnant women were determined.

Prior to inclusion, all potential donors were screened for the inclusion and exclusion criteria and given a detailed instruction sheet with all relevant information with respect to odor donation. Only those living in a smoke-free household were included in the experiment. The donors were provided with detailed instructions such as the avoidance of alcohol, heavily spiced food, garlic and asparagus, which could influence their body odor, at least two days before odor donation. The nights prior to odor donation, the donors were not allowed to share the bed with anyone. These restrictions were meant to prevent the donors’ body odor from being influenced by other scents. The donors were provided with two tight cotton T-shirts (American Apparel), with cellulose pads (Nuk Extra Dry Nursing Pads) sewn into the armpit areas, to be worn during the two nights of odor donation. They also received a pair of pre-washed (with scent-free washing powder) bed linen and towel. Before going to bed, the donors were to use smell-free soap, shampoo and body lotion, all of which had been provided to them. They were instructed not to use other body lotions, perfumes, deodorants or odoriferous substances. The morning following the night-long odor extraction, the T-shirts with the integrated cellulose pads were promptly put into resealable plastic bags and brought to the university hospital by the participants. The pads were separated from the T-shirts and were inspected for traces of deodorant, lotion, or any other scents unrelated to endogenous body odor. The pads were then deep-frozen at -80° Celsius in 1-L freezer bags (Toppits), to be thawed later one hour prior to the fMRI experiment. After odor donation, all donors received a checklist in which they were asked to indicate whether there were any violations of the instructions.

Phase 2: Male Testing

The male participants were recruited at the RWTH Aachen University as well as at the University Hospital in Aachen. Prior to inclusion in the study, the participants had to provide information on sexual orientation, relationship, health status, medication and substance use, and household environment (whether or not it was smoke-free). At this point, the purpose of the study was not revealed to the male participants, who had been informed that some trials would involve exposure to low doses of common odor with other trials involving no odor exposure. The title of the study made available to them read: “Rating of Female Attractiveness”, which described a part of the actual task they would be required to perform. 30 mins prior to the fMRI experiment, 3 odor pads (ovulation odor (OV), pregnant odor (PRG), empty (no-odor) pad (NO)) were thawed using cellulose teabags, with each pad being prepared and used for only one participant. Upon successful completion of the tests, the participants were debriefed.

**Visual stimulation and olfactory stimulation**

The portraits in the Oslo Face database contain a validated attractiveness rating by a Norwegian male sample (n = 20) for each individual picture. 30 photos of women aged 18-30 years, rated as averagely attractive (average attractiveness being defined by a rating between 4.49 and 5.45 on an attractiveness rating scale from 0 (not attractive at all) to 10 (very attractive) (48)) were selected for the experiment.

During the fMRI tasks, the participants were instructed to inhale through the nose at the end of a countdown when a face was presented. Because sniffing in general can initiate movements, the participants were required to practice sniffing without moving the head prior to the actual fMRI experiment. To ensure that the participants would sniff during the presentation of the picture and not during the countdown leading to the image presentation, a Biopac Respiration Belt Transducer was used (https://www.biopac.com/knowledge-base/respiration-recording/).

### **SI fMRI Results**

##### *Effects of task on whole-brain activation*

Investigating different neural activation patterns during attractiveness rating compared to pregnancy categorization and vice versa, independent of condition, we observed activation in several significant clusters. Please see Table S1 for detailed information and Figure S1 for visualization.

Table S1

Brain regions recruited in response to the attractiveness rating > pregnancy categorization (p < .05, FWE-correction) and in response to pregnancy categorization > attractiveness rating (p < .001, extend threshold k = 67, corresponding to Monte Carlo correction)

|  |  |  | Peak voxel | | | |
| --- | --- | --- | --- | --- | --- | --- |
| Anatomical region | Side | k | T | X | Y | Z |
| Attractiveness rating > Pregnancy categorization |  |  |  |  |  |  |
| Middle occipital gyrus  Middle occipital gyrus  Precuneus  Superior parietal lobule  Superior frontal gyrus  Precuneus | R  L  R  R  L  R | 8184 | 10.95  9.60  9.54  9.51  9.27  9.81 | 30  -30  12  16  22  -8 | -70  -88  -62  -64  6  -58 | 36  16  54  54  62  58 |
| Superior frontal gyrus  Precentral gyrus | R | 463 | 9.27  6.09 | 22  34 | 6  -10 | 62  62 |
| Superior frontal gyrus  Middle frontal gyrus | L | 354 | 8.24  8.14 | -26  -24 | 2  2 | 66  62 |
| Precentral gyrus  Inferior frontal gyrus (p. opercularis) | R | 194 | 7.60  7.22 | 44  56 | 8  12 | 30  26 |
| Precentral gyrus  Postcentral gyrus | L | 189 | 7.03  6.82 | -42  -36 | -14  -22 | 64  52 |
| Middle temporal gyrus  Middle occipital gyrus | L | 117 | 7.11  6.92 | -46  -50 | -70  -70 | 6  -25 |
| Precuneus  Cuneus | R | 108 | 7.40  7.18 | 16  20 | -56  -56 | 18  20 |
| Postcentral gyrus | R | 71 | 6.83 | 62 | -16 | 30 |
| Cerebellar vermis | R | 66 | 6.40 | 4 | -44 | -32 |
| Supramarginal gyrus | L | 27 | 6.12 | -52 | -24 | 34 |
| Rolandic operculum | R | 15 | 5.91 | 54 | 10 | -2 |
| Midcingulate gyrus | R | 4 | 5.83 | 8 | 8 | 44 |
| Cerebellum (lobule 6) | L | 4 | 5.59 | -6 | -70 | -20 |
| Pregnancy categorization > Attractiveness rating |  |  |  |  |  |  |
| Cuneus  Superior occipital gyrus | R | 121 | 4.12  4.02 | 10  16 | -92  -90 | 18  20 |

Note. R: right hemisphere; L: left hemisphere

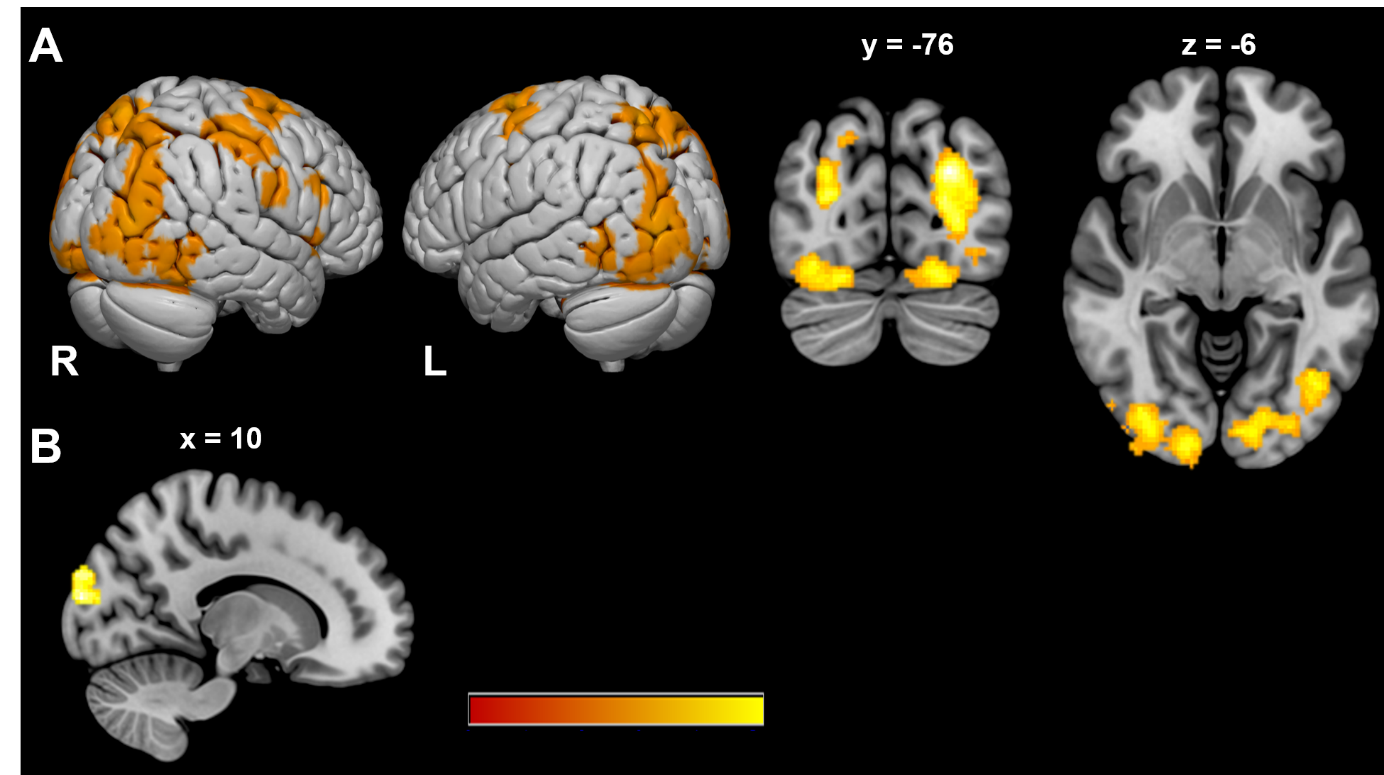

Figure S1. A) Brain regions activated during attractiveness rating > pregnancy categorization. Activation is depicted at p < .05 FWE corrected at voxel level. B) Brain regions activated during pregnancy categorization > attractiveness rating. Significant clusters at p < .001, with extent threshold = 67 (corresponding to Monte Carlo correction).

##### *Effects of body odor on whole-brain activation*

In a whole brain analysis, we evaluated whether body odor in general increases neural activation as compared to NO. While the body odor > NO contrast (both, across tasks and for each task individually) yielded no significant suprathreshold activation in the brain, the opposite contrast (NO > body odor (both, across tasks and for each task individually)) was found to be linked to a widespread activation pattern. Please see Table S2 for detailed information.

Table S2

Brain regions recruited in response to no odor (NO) versus body odor, independent of task and for both tasks individually

|  |  |  | Peak voxel | | | |
| --- | --- | --- | --- | --- | --- | --- |
| Anatomical region | Side | k | T | X | Y | Z |
| NO > Body odor (OV and PRG) |  |  |  |  |  |  |
| Superior frontal gyrus  Posterior-medial frontal gyrus  Posterior-medial frontal gyrus | R  L  R | 519 | 4.86  4.59  4.48 | 18  2  10 | 8  6  16 | 56  48  60 |
| Insula  Inferior frontal gyrus (p. triangularis) | L | 454 | 5.47  4.30 | .40  -36 | 22  22 | 0  10 |
| Rolandic operculum  Inferior frontal gyrus (p. opercularis) | R | 296 | 5.49  4.54 | 59  50 | 4  8 | 10  6 |
| Medial orbital gyrus  Medial orbital gyrus  Anterior cingulate cortex | R  L  R | 296 | 4.61  4.51  3.54 | 10  -2  4 | 52  58  36 | -8  -2  -2 |
| Thalamus  Thalamus | R  L | 251 | 4.27  4.16 | 4  -4 | -14  -12 | 12  12 |
| Posterior-medial frontal gyrus | R | 217 | 5.61 | 0 | -18 | 74 |
| Insula  Putamen  Inferior frontal gyrus (p. triangularis) | R | 213 | 4.58  3.82  3.25 | 36  30  40 | 22  12  24 | 0  8  10 |
| Anterior cingulate cortex  Anterior cingulate cortex  Midcingulate cortex  Superior medial gyrus | L  R  R  L | 188 | 4.42  3.81  3.80  3.50 | -4  4  12  -4 | 30  30  18  52 | 20  24  32  18 |
| Middle frontal gyrus  Inferior frontal gyrus (p. triangularis) | R | 141 | 4.40  3.96 | 38  42 | 42  30 | 32  26 |
| Posterior-medial frontal gyrus  Superior frontal gyrus | L | 108 | 4.23  4.15 | -12  -14 | -2  8 | 74  68 |
| Middle frontal gyrus  Superior frontal gyrus | R | 108 | 4.42  4.17 | 28  23 | 40  30 | 46  56 |
| Middle temporal gyrus  Superior temporal gyrus | R | 108 | 5.25  4.20 | 54  58 | -42  -32 | 4  4 |
| Superior parietal lobule | R | 107 | 4.33 | 26 | -68 | 62 |
| Superior frontal gyrus  Superior medial gyrus | R | 78 | 4.71  4.15 | 16  12 | 62  64 | 26  26 |
| Middle frontal gyrus | L | 77 | 4.22 | -42 | 38 | 32 |
| Middle frontal gyrus | L | 70 | 5.09 | -24 | 58 | 32 |
| Supramarginal gyrus | R | 68 | 4.39 | 60 | -20 | 26 |
| NO > Body odor (OV and PRG) Attractiveness rating |  |  |  |  |  |  |
| Posterior-medial frontal gyrus  Posterior-medial frontal gyrus  Midcingulate cortex  Superior frontal gyrus  Paracentral lobule  Anterior cingulate cortex  Anterior cingulate cortex  Superior medial gyrus | L  R  R  R  L  R  L  R | 2473 | 6.20  5.92  5.85  5.67  5.23  5.19  4.87  4.55 | 0  4  6  26  0  4  0  4 | 8  2  6  -12  -14  32  30  64 | 48  60  44  60  72  22  20  18 |
| Rolandic operculum  Inferior frontal gyrus (p. opercularis)  Insula  Inferior frontal gyrus (p. orbitalis) | R | 1168 | 5.20  5.04  4.93  4.83 | 50  44  36  54 | 6  14  22  22 | 10  6  4  -6 |
| Insula  Inferior frontal gyrus (p. triangularis)  Inferior frontal gyrus (p. opercularis) | L | 977 | 5.67  5.60  5.35 | -42  -32  -46 | 18  30  14 | 2  0  4 |
| Middle frontal gyrus  Inferior frontal gyrus (p. triangularis) | R | 646 | 5.71  4.81 | 30  42 | 38  32 | 30  28 |
| Postcentral gyrus  Supramarginal gyrus | R | 569 | 6.21  5.94 | 60  64 | -18  -18 | 30  28 |
| Middle frontal gyrus  Inferior frontal gyrus (p. triangularis) | L | 456 | 5.55  4.30 | -42  -40 | 36  24 | 30  24 |
| Thalamus  Pallidum  Thalamus | R  L  L | 414 | 4.83  4.58  4.32 | 6  -14  -4 | -20  4  -10 | 12  2  12 |
| Supramarginal gyrus  Superior temporal gyrus | L | 385 | 5.67  5.63 | -64  -64 | -42  -40 | 26  22 |
| Superior frontal gyrus  Posterior-medial frontal gyrus  Precentral gyrus | L | 288 | 4.75  4.35  3.64 | -26  -12  -28 | -6  -2  -6 | 68  74  56 |
| Medial orbital gyrus  Superior orbital gyrus  Medial orbital gyrus  Middle orbital gyrus  Anterior cingulate gyrus | R  R  L  R  R | 280 | 4.71  4.48  4.03  3.67  3.60 | 12  20  -2  24  4 | 48  58  58  52  40 | -6  -2  -8  -4  -4 |
| Precuneus  Superior parietal lobule | L | 278 | 5.03  3.95 | -12  -18 | -52  -66 | 70  64 |
| Middle frontal gyrus  Superior frontal gyrus  Inferior frontal gyrus (p. triangularis) | R | 272 | 4.91  4.03  3.98 | 40  50  28 | 58  62  44 | 0  6  0 |
| Middle temporal gyrus | R | 138 | 4.95 | 54 | -42 | 4 |
| Superior parietal lobule | R | 120 | 4.30 | 20 | -68 | 66 |
| Amygdala | R | 111 | 4.84 | 22 | 6 | -18 |
| Postcentral gyrus  Inferior parietal lobule | L | 110 | 4.74 | -36  -42 | -36  -38 | 44  40 |
| Superior occipital gyrus  Precenues | L | 86 | 4.07  3.56 | -16  -2 | -86  -82 | 48  44 |
| Precentral gyrus | L | 86 | 4.90 | -44 | 2 | 30 |
| Postcentral gyrus  Supramarginal gyrus | L | 74 | 4.91  3.93 | 32  42 | -34  -32 | 40  40 |

Note. R: right hemisphere; L: left hemisphere

Lastly, we sought to determine whether the neural activation during OV or PRG exceeded that during NO, and vice versa. Neither OV > NO nor PRG > NO resulted in significant clusters of activation (both, across tasks and for each task individually). The opposite contrasts (NO > OV and NO > PRG (both, across tasks and for each task individually) resulted in significant activation patterns (Table S3).

Table S3

Brain regions recruited in response to no odor (NO) versus ovulation (OV) and pregnancy odor (PRG) separately, independent of task and for both tasks individually

|  |  |  | Peak voxel | | | |
| --- | --- | --- | --- | --- | --- | --- |
| Anatomical region | **Side** | **k** | **T** | **X** | **Y** | **Z** |
| NO > OV |  |  |  |  |  |  |
| Insula  Inferior frontal gyrus (p. triangularis)  Inferior frontal gyrus (p. orbitalis)  Inferior frontal gyrus (p. opercularis) | L | 242 | 4.89  3.74  3.72  3.33 | -32  -36  -32  -48 | 28  34  33  14 | 4  2  -14  2 |
| Insula  Putamen  Inferior frontal gyrus (p. opercularis)  Inferior frontal gyrus (p. triangularis) | R | 215 | 4.61  3.77  3.60  3.41 | 36  30  40  34 | 26  12  16  30 | -6  6  8  10 |
| Posterior-medial frontal gyrus  Posterior-medial frontal gyrus | R  L | 138 | 4.01  3.82 | 2  -8 | -2  -2 | 58  64 |
| Anterior cingulate gyrus  Superior medial gyrus  Superior medial gyrus | L  R  L | 136 | 4.06  3.91  3.91 | -2  2  2 | 44  60  64 | 16  16  16 |
| Posterior-medial frontal gyrus  Superior frontal gyurs  Posterior medial frontal gyrus | R  L  L | 80 | 3.98  3.79  3.66 | 8  -14  -6 | 16  4  12 | 66  68  70 |
| Inferior frontal gyrus (p. opercularis)  Rolandic Operculum | R | 80 | 4.39  4.06 | 52  50 | 6  4 | 22  10 |
| NO > OV Attractiveness rating |  |  |  |  |  |  |
| Midcingulate cortex  Posterior-medial frontal gyrus  Posterior-medial frontal gyrus  Superior frontal gyrus | R  R  L  R | 1060 | 5.41  5.10  5.09  4.84 | 10  4  0  26 | 18  2  0  -12 | 30  60  60  60 |
| Inferior frontal gyrus (p. opercularis)  Insula  Inferior frontal gyrus (p. triangularis)  Rolandic operculum | R | 662 | 5.41  4.85  4.19  4.10 | 52  36  42  48 | 10  22  18  6 | 18  4  4  10 |
| Postcentral gyrus  Supramarginal gyrus  Rolandic operculum | R | 486 | 5.11  4.95  3.51 | 60  60  68 | -18  -36  -18 | 32  36  16 |
| Insula  Inferior frontal gyrus (p. opercularis)  Inferior frontal gyrus (p. triangularis) | L | 476 | 5.04  4.75  4.69 | -42  -48  -46 | 16  14  18 | 2  2  4 |
| Middle frontal gyrus | R | 441 | 30 | 38 | 30 |  |
| Anterior cingulate cortex  Anterior cingulate cortex | L  R | 190 | 4.37  3.92 | 0  2 | 32  36 | 20  20 |
| Thalamus  Thalamus | R  L | 154 | 4.27  4.06 | 6  -6 | -24  -24 | 12  14 |
| Middle frontal gyrus | L | 78 | 4.16 | -40 | 34 | 30 |
| NO > OV Pregnancy categorization |  |  |  |  |  |  |
| Superior frontal gyrus | R | 122 | 5.18 | 26 | 26 | 58 |
| Paracentral lobule | L | 71 | 4.48 | -2 | -20 | 76 |
| NO > PRG |  |  |  |  |  |  |
| Insula  Inferior frontal gyrus (p. triangularis) | L | 414 | 5.21  4.52 | -40  -34 | 22  30 | 0  4 |
| Middle orbital gyrus  Middle orbital gyrus  Anterior cingulate gyrus  Superior orbital gyrus  Superior frontal gyrus | L  R  R  R  R | 373 | 4.60  4.54  4.16  4.13  3.88 | -2  12  4  20  22 | 58  48  36  58  56 | -2  -6  -2  -2  0 |
| Rolandic operculum  Inferior frontal gyrus (p. opercularis)  Inferior frontal gyrus (p. orbitalis) | R | 328 | 5.39  4.53  3.65 | 50  50  54 | 4  8  22 | 10  6  -4 |
| Posterior-medial frontal gyrus  Posterior-medial frontal gyrus  Midcingulate cortex | L  R  R | 319 | 5.13  3.64  3.61 | -4  4  2 | 10  2  -8 | 46  58  50 |
| Thalamus  Thalamus | L  R | 210 | 4.49  4.33 | -4  12 | -8  -20 | 10  10 |
| Paracentral lobule | L | 183 | 5.00 | -2 | -16 | 74 |
| Middle frontal gyrus  Inferior frontal gyrus (p. triangularis)  Superior frontal gyrus | R | 164 | 4.56  4.32  3.89 | 30  42  26 | 36  30  38 | 46  26  48 |
| Middle frontal gyrus | L | 144 | 4.71 | -42 | 40 | 28 |
| Olfactory cortex  Insula | R | 130 | 4.63  3.39 | 22  12 | -18  12 | -18  -16 |
| Supramarginal gyrus | L | 119 | 4.09 | -62 | -42 | 26 |
| Superior medial gyrus  Middle frontal gyrus | R | 115 | 4.90  3.52 | 12  24 | 64  58 | 26  20 |
| Putamen | L | 101 | 4.72 | -20 | 4 | -10 |
| Superior frontal gyrus | L | 80 | 4.52 | -26 | -10 | 70 |
| Middle temporal gyrus | L | 79 | 4.42 | -44 | -66 | 14 |
| Posterior-medial frontal gyrus  Superior frontal gyrus | R | 78 | 4.24  4.29 | 10  18 | 16  8 | 60  56 |
| Middle temporal gyrus | R | 70 | 4.52 | 56 | -42 | 4 |
| NO > PRG Attractiveness rating |  |  |  |  |  |  |
| Midcingulate cortex  Posterior-medial frontal gyrus  Posterior-medial frontal gyrus  Anterior cingulate cortex | R  L  R  R | 1882 | 6.46  6.00  5.16  4.72 | 6  2  4  6 | 6  6  2  32 | 44  48  58  22 |
| Supramarginal gyrus  Superior temporal gyrus  Middle temporal gyrus  Heschl’s gyrus  Angular gyrus  Superior temporal gyrus | L | 842 | 6.80  6.64  4.89  4.46  4.26  4.15 | -64  -64  -64  -48  -56  -52 | -42  -40  -32  -16  -58  -40 | 26  22  8  6  32  20 |
| Rolandic operculum  Inferior frontal gyrus (p. orbitalis)  Inferior frontal gyrus (p. opercularis)  Temporal pole | R | 670 | 5.45  5.15  4.44  3.97 | 56  56  44  46 | -2  22  14  22 | 6  -4  6  -18 |
| Insula  Inferior frontal gyrus (p. triangularis)  Inferior frontal gyrus (p. opercularis) | L | 606 | 5.31  5.20  4.38 | -34  -32  -46 | 24  30  14 | -2  0  4 |
| Middle frontal gyrus  Inferior frontal gyrus (p. triangularis) | L | 578 | 5.84  4.39 | -42  -40 | 40  24 | 28  24 |
| Superior frontal gyrus  Middle orbital gyrus  Middle orbital gyrus  Middle frontal gyrus | R  R  L  R | 570 | 4.97  4.67  4.62  4.41 | 20  12  -2  40 | 58  48  58  58 | 0  -6  -2  0 |
| Postcentral gyrus  Superior temporal gyrus  Supramarginal gyrus  Rolandic operculum | R | 386 | 5.53  5.35  4.76  3.72 | 60  70  60  54 | -18  -28  -32  -16 | 30  18  30  18 |
| Inferior frontal gyrus (p. triangularis)  Middle frontal gyrus | R | 386 | 4.64  4.49 | 44  30 | 36  40 | 26  28 |
| Thalamus  Thalamus | R  L | 313 | 4.92  4.65 | 8  -4 | -16  -10 | 10  12 |
| Calcarine gyrus  Calcarine gyrus | L  R | 298 | 4.50  4.45 | -2  2 | -74  -74 | 16  12 |
| Amygdala  Insula | R | 214 | 4.84  4.40 | 24  28 | 6  18 | -16  -14 |
| Precunues  Superior parietal lobule | L | 208 | 5.21  4.19 | -12  -18 | -52  -66 | 70  66 |
| Superior medial gyrus  Superior medial gyrus | R  L | 159 | 5.00  3.63 | 6  -6 | 64  68 | 20  14 |
| Middle temporal gyrus  Superior temporal gyrus | R | 134 | 4.85  3.78 | 58  68 | -34  -24 | 4  4 |
| Postcentral gyrus  Inferior parietal lobule | L | 121 | 4.91  4.19 | -36  -46 | -36  -28 | 44  42 |
| Superior occipital gyrus  Precunues | L | 116 | 3.96  3.83 | -16  -8 | -86  -82 | 48  48 |
| Middle frontal gyrus  Middle orbital gyrus  Inferior frontal gyrus (p. triangularis) | L | 105 | 4.08  4.01  3.84 | -36  -36  -36 | 56  50  42 | 8  -2  0 |
| Superior parietal lobule | R | 82 | 4.63 | 20 | -68 | 66 |
| Anterior cingulate cortex  Anterior cingulate cortex | L  R | 67 | 4.04  3.74 | 2  2 | 46  42 | 16  18 |

Note. R: right hemisphere; L: left hemisphere
